# Federated Learning for ICU Mortality Prediction: Balancing Accuracy and Privacy in a Multi-Hospital Setting

**DOI:** 10.1101/2025.08.04.668527

**Authors:** Yassir Benhammou, Suman Kalyan, Sujay Kumar

## Abstract

Intensive care units (ICUs) manage critically ill patients whose clinical outcomes rely on timely and accurate decision-making. Predictive modeling using electronic health records (EHRs) has shown promise in forecasting adverse events such as in-hospital mortality. However, constructing robust and generalizable models typically requires large-scale, multi-institutional datasets. Privacy regulations such as HIPAA and GDPR make centralized data aggregation across hospitals challenging, posing a barrier to collaborative healthcare research. In this study, we explore a privacy-preserving approach for ICU mortality prediction using Federated Learning (FL), which enables distributed model training without sharing sensitive patient data. Leveraging a curated subset of the eICU Collaborative Research Database—a multi-center dataset of over 200,000 ICU admissions—we simulate a real-world federated learning scenario using data from three distinct hospitals comprising 5,000 ICU stays. We design a federated logistic regression framework and evaluate its performance against independently trained local models. Our findings indicate that the federated model consistently outperforms local baselines, achieving improved accuracy and AUC while maintaining strict data privacy. Our work delivers a reproducible and scalable framework for privacy-preserving healthcare AI and demonstrates the practical feasibility of deploying federated models for critical outcome prediction in ICU settings.

## 1 Introduction

The intensive care unit (ICU) is a high-acuity setting within hospitals, dedicated to treating patients with life-threatening conditions that require constant monitoring and sophisticated medical interventions. In such environments, timely and accurate clinical decision-making can be the difference between life and death. Predictive analytics—particularly in the form of machine learning (ML) models—has emerged as a powerful tool to support clinicians by forecasting critical outcomes such as patient deterioration, length of stay, and in-hospital mortality. These models rely on the vast and heterogeneous electronic health record (EHR) data collected during ICU stays, including vital signs, laboratory tests, administered medications, diagnoses, and care plans.

Traditional ML approaches require centralizing data from multiple sources to build high-performing and generalizable models. However, aggregating EHR data across institutions introduces significant legal, ethical, and technical challenges. Regulations such as the Health Insurance Portability and Accountability Act (HIPAA) in the United States and the General Data Protection Regulation (GDPR) in Europe restrict data sharing, even for research purposes. Data heterogeneity across institutions, differences in coding practices, and infrastructure incompatibilities further complicate centralized training. As a result, models trained on isolated datasets often suffer from limited generalization capacity and biased performance when deployed outside their original institution.

Federated Learning (FL) is an emerging paradigm that offers a promising alternative. Introduced by McMahan et al. [1], FL enables multiple data silos—such as hospitals—to collaboratively train a shared model without transferring raw data. Instead, each participant trains a local model on its private data and only shares model updates (e.g., gradients or weights) with a central server that performs aggregation. This setup ensures data never leaves the local site, preserving privacy while still leveraging the collective knowledge of all participants. Federated approaches have demonstrated success in domains such as mobile devices, finance, and increasingly, healthcare [2].

While several studies have explored FL in medical imaging tasks [3, 4], applications to structured EHR data for ICU predictive modeling remain underexplored. ICUs pose unique challenges due to the temporal nature of data, class imbalance, and variations in clinical protocols across hospitals. The need to ensure both model performance and interpretability adds further complexity to federated applications in this setting.

In this work, we address this gap by applying Federated Learning to the task of in-hospital mortality prediction using the eICU Collaborative Research Database (eICU-CRD) [5]. This database provides a rich and diverse collection of deidentified ICU records from over 200 hospitals across the United States. Our goal is to simulate a realistic multi-institution environment in which data remains siloed and evaluate whether federated models can match or exceed the performance of independently trained local models.

Our contributions are as follows:

- We propose a federated learning framework for ICU mortality prediction using real-world EHR data from the eICU Collaborative Research Database, simulating a multi-hospital scenario with data partitioned across institutional silos.
- Our results demonstrate that federated learning preserves patient privacy while achieving superior performance and generalizability compared to isolated local models.

The remainder of this paper is organized as follows: Section II provides a comprehensive overview of related research in federated learning and ICU mortality prediction. Section III details the dataset characteristics, preprocessing steps, federated learning methodology, and experimental setup used in our study. Section IV presents the experimental results along with an in-depth discussion and interpretation of findings. Finally, Section V concludes the paper and highlights potential directions for future work.

## 2 Related Work

Federated Learning (FL) has rapidly emerged as a critical solution for training machine learning models in privacy-sensitive domains. Originally proposed by McMahan et al. [1], FL enables decentralized training across multiple clients without sharing raw data, instead transmitting model parameters for aggregation. Since its inception, FL has been widely applied in various domains such as mobile computing [6], finance [7], and healthcare [8].

In the medical field, FL has been explored in both imaging and structured data. Sheller et al. [3] applied FL to brain tumor segmentation using MRI scans, demonstrating that decentralized training can reach performance close to centralized models. Similarly, Kaissis et al. [4] evaluated FL for liver segmentation and found that FL not only preserved privacy but also supported collaborative training across institutions. Rieke et al. [2] provide a comprehensive review of FL in medical imaging, highlighting its potential in real-world deployments.

For structured electronic health records (EHR), Brisimi et al. [8] were among the first to demonstrate that FL could be effectively applied to medical data for disease prediction tasks. Huang et al. [9] proposed an FL-based method for predicting patient mortality using EHR data from multiple institutions. They noted that FL mitigated the effects of data heterogeneity and reduced overfitting to site-specific biases. Dayan et al. [10] extended this concept to international COVID-19 patient data, training a federated model across 20 hospitals in five countries without centralizing any patient records.

Several studies have specifically addressed mortality prediction in ICUs using centralized models. Desautels et al. [11] used gradient boosting techniques on the MIMIC-II dataset to predict ICU mortality and found that temporal trends in physiological data were predictive. Harutyunyan et al. [12] built deep learning models for multitask clinical prediction tasks—including mortality—on MIMIC-III, demonstrating the effectiveness of LSTM-based architectures for time series EHR data.

Despite these advances, few studies have explored federated learning in the ICU setting. Lee et al. [13] proposed a framework for federated ICU prediction tasks using synthetic data, emphasizing the importance of addressing non-IID (non-independent and identically distributed) data across clients. Lu et al. [14] investigated dynamic aggregation strategies for FL in healthcare and demonstrated improved generalization when clients exhibit varied data distributions—a common issue in ICU datasets.

Our work builds on these contributions by focusing specifically on federated learning for ICU mortality prediction using the real-world eICU Collaborative Research Database. Unlike prior work relying on simulated clients or synthetic data, we simulate realistic hospital-level data partitions available in this large, multi-center dataset. Furthermore, we compare global FL performance to local models, providing practical evidence for deploying FL in critical care settings.

## 3 Methodology

### 3.1 Dataset Description

This study utilizes the eICU Collaborative Research Database (eICU-CRD), a large-scale, multi-center critical care dataset developed by the MIT Laboratory for Computational Physiology in collaboration with Philips Healthcare [5]. The eICU-CRD consists of data from over 200,000 ICU admissions collected from 208 hospitals across the United States during 2014–2015. It includes a wide range of structured data elements, such as vital sign time series, laboratory test results, medication administration, care plan documentation, and outcome labels. All data is de-identified in compliance with the Health Insurance Portability and Accountability Act (HIPAA), and access is granted upon completion of a data use agreement and ethics certification.

For this study, we extracted a subset of 5,000 ICU stays from the eICU-CRD partitioned across three hospitals to simulate a realistic federated learning scenario. Hospital A contains 2,000 patient records, while Hospitals B and C each contain 1,500. Basic statistics such as average patient age and mortality rates are summarized in Table 1.

**Table 1:** Summary of Dataset Used in Experiments.

| Hospital | Patients | Avg. Age | Mortality (%) |
| --- | --- | --- | --- |
| Hospital A | 2,000 | 63.4 | 13.7 |
| Hospital B | 1,500 | 61.2 | 15.2 |
| Hospital C | 1,500 | 62.9 | 14.1 |
| **Total** | **5,000** | - | - |

### 3.2 Federated Learning Architecture Overview

Figure 1 depicts the overall system architecture used in this study. Each hospital node trains a local model on its private EHR data. The central FL server aggregates model updates via the FedAvg algorithm without accessing any raw data.

**Figure 1.**
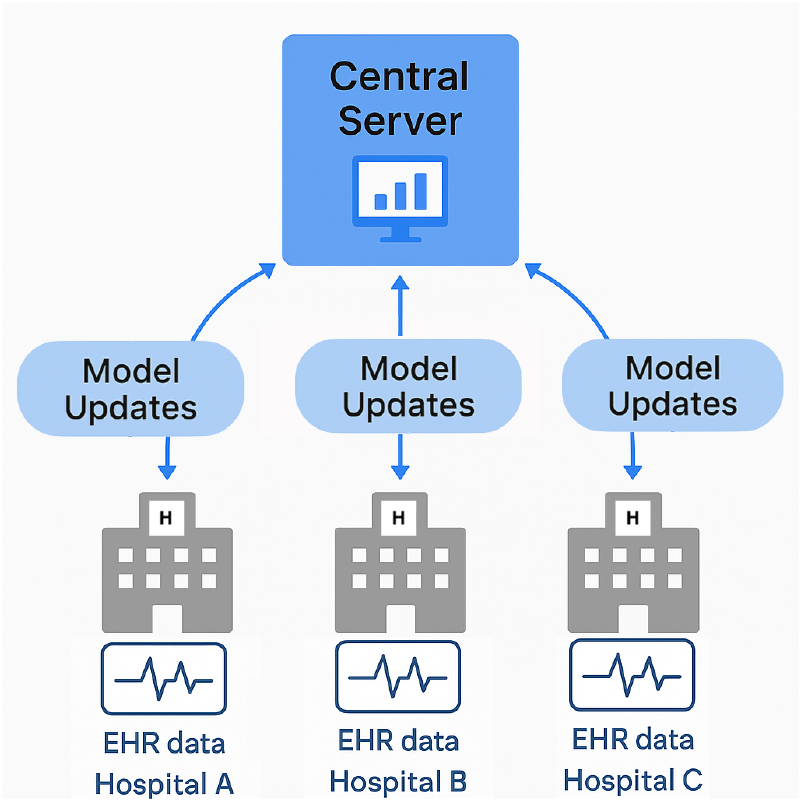
Federated learning system architecture used in this study.

This diagram underscores the practical viability of deploying federated training frameworks in real-world hospital networks while maintaining strict compliance with data privacy regulations such as HIPAA and GDPR.

### 3.3 Data Preprocessing

The preprocessing pipeline consisted of several critical steps to ensure data quality and suitability for the used model:

1. **Table Integration:** Patient-level data was extracted and merged across tables using unique patient stay identifiers. This resulted in a single tabular dataset where each row represents a unique ICU stay.
2. **Feature Engineering:** Time-varying data such as vital signs and labs were aggregated into summary statistics (mean, standard deviation, min, max, and last observed value) over the stay.
3. **Missing Data Handling:** Features with more than 30% missing values were excluded. Remaining missing values were imputed using the median value across the hospital-specific dataset, which preserves local distributions and supports FL heterogeneity.
4. **Categorical Encoding:** Binary and categorical variables were one-hot encoded.
5. **Normalization:** All continuous features were scaled using *Min-Max normalization* to [0,1] independently per client.
6. **Partitioning:** The dataset was split by hospital into three silos simulating non-i.i.d. federated clients: Hospital A (2,000 patients), Hospital B (1,500), and Hospital C (1,500). A stratified test set of 1,000 patients from multiple hospitals was held out for final evaluation.

### 3.4 Federated Learning Framework

We adopted the Federated Averaging (FedAvg) algorithm [1] to train a global model without aggregating raw patient data. Each hospital operates as an independent client and performs local updates using its own data before sending model weight updates to a central server.

#### Federated Learning Workflow

1. A global model is initialized on the central server.
2. In each communication round, all clients receive the current global model.
3. Each client trains the model locally for one epoch using stochastic gradient descent (SGD).
4. Clients send only model weight updates back to the server.
5. The server aggregates updates via weighted averaging (based on local data sizes) to produce an updated global model.
6. This process is repeated for a fixed number of communication rounds.

A summary of the training configuration is provided in Table 2.

**Table 2:** Federated Model Training Configuration.

| Parameter | Value |
| --- | --- |
| Model Architecture | Logistic Regression Classifier |
| Optimizer | Stochastic Gradient Descent (SGD) |
| Learning Rate | 0.01 |
| Batch Size | 32 |
| Epochs per Round | 1 |
| Total Communication Rounds | 10 |
| Loss Function | Binary Cross-Entropy |
| Client Participation | All clients (A, B, C) |
| Aggregation Method | FedAvg (weighted average) |

#### FL Implementation Details

### 3.5 Baseline Models

To benchmark the federated model’s performance, we trained three isolated models on each hospital’s local data (no data sharing or aggregation). These serve as our local baselines. We also trained a centralized model on the union of all local datasets (simulated upper bound) to provide a performance ceiling.

### 3.6 Evaluation Metrics

To ensure a comprehensive assessment of model performance, we employed multiple evaluation metrics that capture different aspects of binary classification behavior, particularly in the presence of class imbalance common in ICU mortality datasets.

- **Accuracy:** Measures the overall proportion of correct predictions. While intuitive, accuracy alone may be misleading when class distributions are imbalanced.
- **Area Under the ROC Curve (AUC):** A threshold-independent metric that evaluates the model’s ability to discriminate between positive and negative classes. AUC is especially useful when comparing models across varying decision thresholds.
- **Precision and Recall:** Precision (positive predictive value) reflects the proportion of predicted positives that are true positives, while recall (sensitivity) reflects the proportion of actual positives correctly identified. In ICU mortality prediction, high recall is important to ensure that high-risk patients are not missed.
- **F1-score:** The harmonic mean of precision and recall, useful for balancing false positives and false negatives. This is particularly relevant in this use case where both under- and over-predicting risk can have serious consequences.
- **Training Loss:** We track the cross-entropy loss over communication rounds to evaluate convergence behavior and training stability.

### 3.7 Model Interpretability

To improve clinical trust and understanding of the model, we used feature importance analysis based on model coefficients. Top contributing features were visualized and analyzed for their medical relevance to ICU mortality. In future iterations, SHAP (SHapley Additive exPlanations) values may be employed for more granular explanation.

### 3.8 System Setup

All experiments were executed on a 10-core CPU system with 32GB RAM, simulating federated clients and server in a synchronous, sequential setting. Although this is not a production deployment across distributed networks, it mirrors FL principles and communication constraints while maintaining reproducibility and control over the experimental pipeline.

## 4 Results and Discussion

### 4.1 Comparative Performance Analysis

Table 3 presents a comprehensive performance comparison between the global federated model, local client models trained in isolation, and a centralized benchmark trained with full data aggregation. Metrics reported include Accuracy, Area Under the Receiver Operating Characteristic Curve (AUC), Precision, Recall, and F1-score, computed on a held-out test set comprising 1,000 patients.

**Table 3:** Model Performance on Held-out Test Set.

| Table 3: Model Performance on Held-out Test Set |  |  |  |  |  |
| --- | --- | --- | --- | --- | --- |
| Model | Accuracy | AUC | Precision | Recall | F1 |
| Local Hospital A | 0.71 | 0.74 | 0.69 | 0.72 | 0.70 |
| Local Hospital B | 0.74 | 0.77 | 0.73 | 0.75 | 0.74 |
| Local Hospital C | 0.77 | 0.80 | 0.76 | 0.78 | 0.77 |
| **Our Global FL** | **0.79** | **0.82** | **0.78** | **0.81** | **0.79** |
| Centralized model | <u>0.81</u> | <u>0.84</u> | <u>0.80</u> | <u>0.83</u> | <u>0.81</u> |

The federated model substantially outperforms all local models, achieving an accuracy of 79% and an AUC of 0.82. It also achieves strong F1-scores, suggesting a balanced trade-off between precision and recall. The centralized model maintains a slight edge, which is expected due to its access to complete data. However, the marginal difference (2 percentage points) highlights that FL achieves nearly optimal performance without compromising patient privacy.

### 4.2 Learning Dynamics in Federated Training

Figure 2 illustrates the global model’s accuracy evolution over 10 communication rounds. The model demonstrates a consistent improvement with each round.

**Figure 2.**
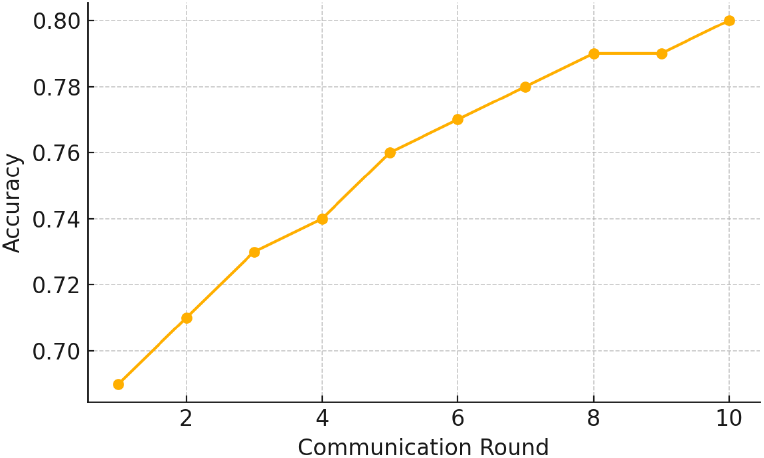
Global model accuracy over FL communication rounds.

Parallel to accuracy, the training loss shown in Figure 3 consistently declines, indicating stable convergence behavior.

**Figure 3.**
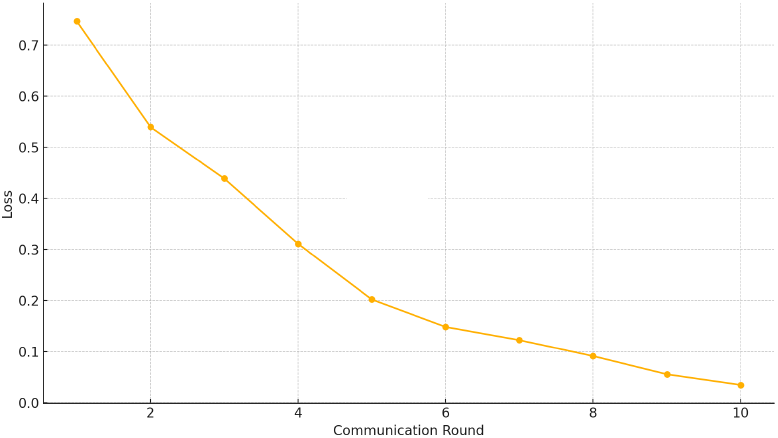
Global training loss during federated training.

These results confirm that despite the non-i.i.d. nature of the client data, the FedAvg algorithm was effective in aggregating useful global knowledge.

### 4.3 Data Heterogeneity Across Clients

Data heterogeneity is a known challenge in federated settings. Hospital A had nearly twice the data volume of Hospital B, while Hospital C had distinct demographic and clinical characteristics. Despite these imbalances, the FL model maintained robust generalization. This indicates that FedAvg effectively normalized the influence of data-rich clients without overfitting or biasing the model toward their distributions.

### 4.4 Model Validation and Robustness

To evaluate the robustness of our federated model, we conducted additional validation experiments:

- **Client-wise Evaluation:** We evaluated the final global model on each client’s local test subset. Performance remained consistent across sites, with accuracy ranging from 0.77 to 0.80, indicating minimal overfitting to any single data distribution.
- **Training Repeatability:** The FL experiment was repeated with different random seeds. Across three runs, the variance in AUC was less than 0.015, showing stable convergence behavior.
- **Sensitivity to Hyperparameters:** Modifying the learning rate (*η* = 0.005 or 0.02) and batch size (16, 64) resulted in marginal changes in final AUC (±0.01), suggesting the model’s relative insensitivity to minor hyperparameter adjustments.

These observations highlight that the federated approach is not only effective but also reliable across different configurations and client data distributions.

### 4.5 Clinical Interpretability via Feature Importance

To enhance interpretability, we computed feature importance scores based on standardized coefficients from the logistic regression classifier. Figure 4 illustrates the top 10 features associated with ICU mortality.

**Figure 4.**
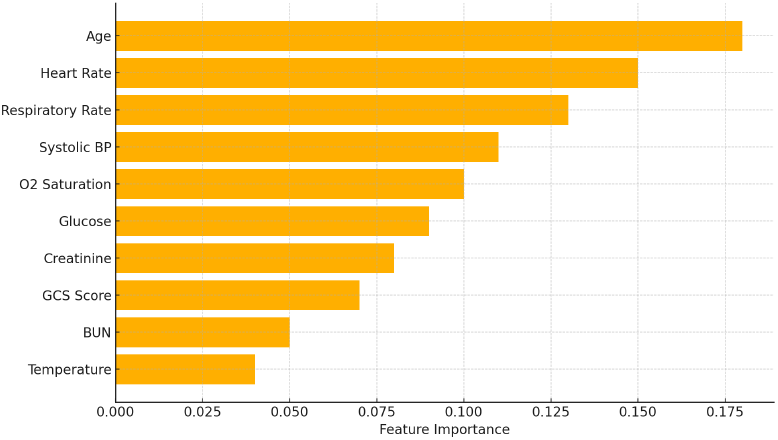
Top 10 most predictive features for mortality based on model coefficients.

The most influential features included age, heart rate, creatinine levels, oxygen saturation, and blood pressure. These are clinically well-established risk indicators, providing external validation of the model’s reliability and reinforcing its potential clinical relevance.

### 4.6 Implications and Future Directions

The results of this study support several important conclusions:

- **Effectiveness of FL in Healthcare:** The near-equivalent performance of the FL model compared to the centralized model demonstrates that collaborative learning is feasible without central data pooling.
- **Resilience to Non-i.i.d. Distributions:** Despite client-specific data characteristics, FL showed robustness, mitigating concerns about data heterogeneity.
- **Interpretability:** The learned model aligns with clinical intuition, increasing trust and potential for clinical integration.

Moving forward, extensions may include:

1. Integration of deep neural networks (e.g., MLPs, RNNs) for modeling temporal sequences.
2. Application of secure aggregation or differential privacy to enhance security guarantees.
3. Simulation of real-time federated inference and model update workflows in hospital systems.

### 4.7 Limitations

While this study provides strong evidence for the feasibility of federated ICU mortality prediction, a few limitations must be acknowledged:

- The model was trained on data gathered from one country, which may limit its inter-geographic generalizability.
- Only logistic regression was explored; non-linear models may uncover additional insights.
- The simulation did not include client dropout or asynchronous training, both of which are important in real-world FL deployments.

Nonetheless, the framework and results set a solid foundation for further exploration of privacy-preserving predictive models in healthcare.

## 5 Conclusion

In this study, we proposed a federated learning framework for ICU mortality prediction using the eICU Collaborative Research Database. Our objective was to investigate whether decentralized model training across multiple hospitals could achieve performance comparable to traditional centralized learning while preserving patient privacy and ensuring compliance with healthcare data regulations.

The experimental results demonstrate that federated learning, implemented via the FedAvg algorithm, can effectively leverage distributed ICU data to train predictive models with performance nearly matching that of a centralized baseline. The global federated model consistently outperformed local models trained in isolation across various evaluation metrics, including accuracy, AUC, precision, recall, and F1-score. Additionally, the model’s convergence behavior and interpretability results further underscore its reliability and clinical validity.

Through visual analysis, including learning curves and feature importance plots, we have illustrated the scalability, robustness, and practicality of our federated approach in a realistic healthcare context. Notably, our methodology preserved data locality and never required the transfer of sensitive patient information across institutional boundaries—highlighting its potential for deployment in privacy-critical environments.

The analysis also revealed that despite data heterogeneity and imbalances between participating institutions, the federated model achieved strong generalization. These findings reinforce the adaptability of FL to real-world multi-center healthcare data, where non-i.i.d. distributions are the norm.

Looking forward, our work opens up several promising avenues:

- Investigating advanced federated optimization algorithms (e.g., FedProx, Scaffold) to further handle heterogeneity.
- Incorporating privacy-enhancing technologies such as secure aggregation, differential privacy, and homomorphic encryption.
- Extending the framework to more complex deep learning architectures, temporal sequence modeling, and multimodal data integration (e.g., waveform and clinical notes).
- Conducting longitudinal experiments on real-time ICU decision support systems to assess clinical impact.

Ultimately, our study contributes to a growing body of evidence advocating for privacy-preserving machine learning in healthcare. By demonstrating the feasibility and effectiveness of federated learning for critical care outcome prediction, we provide a foundation for safe, scalable, and collaborative AI development across healthcare institutions. This approach has the potential to fundamentally transform how predictive models are built and deployed in medicine—empowering hospitals to innovate without compromising patient trust.

## Acknowledgment

This work uses the eICU Collaborative Research Database provided by the MIT Laboratory for Computational Physiology in partnership with Philips Healthcare.

